# Demuxalot: scaled up genetic demultiplexing for single-cell sequencing

**DOI:** 10.1101/2021.05.22.443646

**Authors:** Alex Rogozhnikov, Pavan Ramkumar, Kevan Shah, Rishi Bedi, Saul Kato, G. Sean Escola

## Abstract

Demultiplexing methods have facilitated the widespread use of single-cell RNA sequencing (scRNAseq) experiments by lowering costs and reducing technical variations. Here, we present *demuxalot*: a method for probabilistic genotype inference from aligned reads, with no assumptions about allele ratios and efficient incorporation of prior genotype information from historical experiments in a multi-batch setting. Our method efficiently incorporates additional information across reads originating from the same transcript, enabling up to 3x more calls per read relative to naive approaches. We also propose a novel and highly performant tradeoff between methods that rely on reference genotypes and methods that learn variants from the data, by selecting a small number of highly informative variants that maximize the marginal information with respect to reference single nucleotide variants (SNVs). Our resulting improved SNV-based demultiplex method is up to 3x faster, 3x more data efficient, and achieves significantly more accurate doublet discrimination than previously published methods. This approach renders scRNAseq feasible for the kind of large multi-batch, multi-donor studies that are required to prosecute diseases with heterogeneous genetic backgrounds.

## INTRODUCTION

Single-cell RNA sequencing (scRNAseq) has become cheaper, more standardized and ubiquitous thanks to pooled droplet-based designs (Hinz et al. 2019; Macosko et al. 2015) that have been commercialized by platforms such as 10X Genomics (Zheng et al. 2017). When working with multiple genotypes, as is increasingly common in disease biology, computational demultiplexing methods that recover the genotype identity of single cells offer several advantages. First, by allowing pooling across samples from multiple genotypes in the same run, they increase throughput and decrease the overall cost of sequencing. Second, pooling eliminates lane-specific confounding effects. Third, undetected confounding effects arising from sample index swapping (Griffiths et al. 2018), mishandling, or cell-line identity contamination (Griffiths et al. 2018; Korch et al. 2018) during library prep or sequencing steps can easily be detected and corrected post hoc. Finally, demultiplexing methods are effective at detecting and eliminating doublets, whose likelihood increases with sample pooling due to a higher likelihood of encapsulating multiple cells per droplet. Therefore, there is a strong motivation to improve demultiplexing methods for accuracy, robustness, speed, and data efficiency to support a large number of genotypes.

Existing demultiplexing methods use SNVs to discriminate between genotypes since they explain the largest genetic variability and are most reliably detected by next-generation sequencing (NGS) technologies. Methods differ by whether they use an independent source of genotype information for SNVs (Kang et al. 2018), such as low-cost DNA microarrays (Macdonald et al. 2005; Fan et al. 2006), or whether these are learned directly from the data (Xu et al. 2019; Heaton et al., 2019; Huang, McCarthy, and Stegle 2019). Methods that restrict themselves to commonly available reference SNVs, such as from the 1000 Genomes Project (Consortium and The 1000 Genomes Project Consortium 2015), are limited because they do not exploit additional putative SNVs specific to the donors in the dataset. Therefore, several published methods propose techniques to detect additional putative SNVs from data (Xu et al. 2019; Heaton et al., 2019; Huang, McCarthy, and Stegle 2019). The high number of putative SNVs with low relevance on the other hand makes genotype deconvolution computationally expensive and introduces noise. Therefore, demultiplexing methods could benefit from a principled approach to optimally exploit additional putative SNVs.

Multi-donor experiments are increasingly common and important for characterizing disease biology using complex *in vitro* models, particularly in a scaled setting. Further, in settings where data for different donors are acquired in different experimental batches, it is not uncommon that reference SNVs for different donors are obtained from different genotyping arrays, resulting in heterogeneity of available genetic information. Current demultiplexing methods, not designed to scale beyond a single experiment, would deal with these discrepancies by restricting the data – specifically, by discarding SNVs not shared across donors. Therefore, demultiplexing methods could potentially be designed to better accumulate genotype evidence from historical data.

Demultiplexing quality directly depends on the discriminative power of SNVs used, which in turn depends on total number of SNVs, the number of SNVs called per read, and the confidence with which they are called. Therefore, demultiplexing efficiency can benefit from three improvements: optimally selecting new SNVs using current best estimates of the genotypes to maximize information gain, (2) optimally accumulating prior genotype evidence obtained from different batches with potentially different sources of reference SNVs, and (3) capturing more calls by using multiple reads from the same transcript molecule for deep sequencing settings.

Here, we present a method that improves upon published methods on all these fronts. Our conjugate Bayesian model smoothly integrates genotype information from reference SNVs and dataset-specific detected putative SNVs, as well as from historical experiments in a multi-batch setting. Additionally, we propose improvements for more data efficient SNV calling across reads from a transcript, resulting in up to 3x more calls per read. Finally, we propose computationally efficient selection of putative SNVs, optimizing for differential information gain for the dataset-specific genotypes. Our model makes fewer assumptions about allele ratios at SNV loci, iterates between genotype assignment and genotype refinement steps in an expectation–maximization algorithm, and converges in very few iterations. We show using two different ground truth datasets that our method is faster, more accurate for genotype assignment and doublet detection, and more data efficient than prior methods. We also provide an efficient Python implementation as an open source library (https://github.com/herophilus/demuxalot).

## RESULTS

We developed *demuxalot*, a genotype demultiplexing algorithm for pooled scRNAseq experiments to address the limitations of existing methods in scaling to a large number of donors or batches. In droplet-based designs, individual reads carry unique genotypic information on known SNVs as well as putative SNVs detectable from variation in the dataset. Once sequenced and aligned, reads can be mapped back to their transcript and droplet of origin due to unique molecular indicator (UMI) tags and droplet-specific barcodes respectively. The computational problem of demultiplexing may then be stated as the inference of probabilistic genotype assignments for each droplet from genotypic information available in each read.

Demultiplexing methods employ some set of the following steps: **(1) SNV calling**: all methods begin by calling SNVs at genomic loci from aligned reads; **(2) Base frequency or allele estimation**: called SNVs are somehow aggregated across transcripts from each barcode to estimate base frequencies or allele ratios at each SNV location; **(3) Genotype initialization**: genotype signatures, defined as the estimates of the allele frequencies at each SNV locus for a given genotype, are initialized from an external source (e.g. bead arrays) or from prior batches, or are inferred directly from the data; **(4) Genotype assignment**: genotype identities are assigned to each barcode based on called SNVs and current genotype signature estimates; **(5) Genotype refinement**: genotype signatures are refined based on called SNVs and current genotype assignments; and, in some methods, **(6) SNV enrichment**: as genotype information provided by bead arrays or estimated from prior runs may be incomplete or partially incorrect, novel putative SNVs are detected from non-reference loci on the genome (Xu et al. 2019; Heaton et al., 2019; Huang, McCarthy, and Stegle 2019).

Below, we highlight specific ways in which *demuxalot* improves upon published methods in each of these steps. Briefly, we model the demultiplexing inference problem based on a generative process of the aligned reads (**Fig. 1a**) which permits us to develop a straightforward Bayesian framework for genotype inference (**Fig. 1b–c; see Methods**). Each barcoded droplet comprises several thousand unique transcripts (molecules), each of which are fragmented, amplified and sequenced as reads. We then use aligned reads and an a priori estimate of the genotype signature, obtained either from SNV bead arrays or from historical runs for each known genotype at each SNV, to infer the genotype identity of each barcode. The base frequencies (**α**) are modeled as Bernoulli random variables for each base at each SNV for each droplet barcode, the genotype signatures (**β**) as Dirichlet random variables at each SNV for each genotype, and the genotype assignments (**γ**) as categorical random variables for each barcode. These distributional assumptions not only simplify model estimation, but also enable natural extensions of published methods for scaling to large numbers of donors and batches.

**Figure 1.**
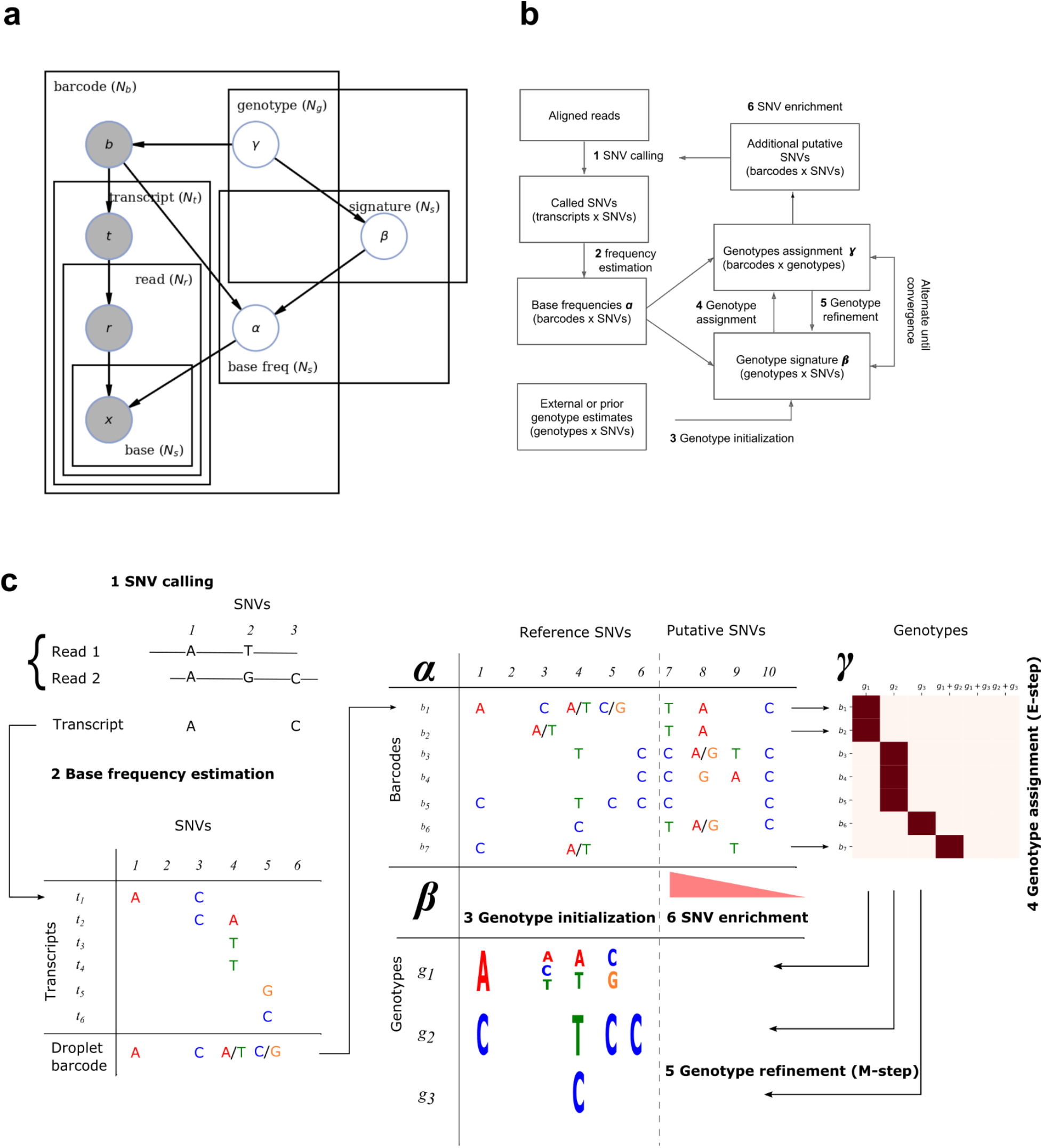
Demuxalot algorithm. (**b**) Generative model of aligned reads from single-cell sequencing methods expressed in plate notation. Observed variables are in filled circles and estimands are in unfilled circles. Droplets tagged with a unique barcode, ***b*** can have cells or multiplets expressing many transcripts, ***t***. Each transcript gets amplified, fragmented into reads ***r***, sequenced and subsequently aligned to the reference genome. A read can overlap multiple SNVs at bases ***x***. The demultiplexing problem can be stated as the problem of simultaneously inferring base frequencies **α** at each reference SNV or putative SNV ***s***, the signature of each genotype **β**, and the probabilistic assignment of each barcode to a genotype **γ**. (**b–c**) Overview (**b**) and detailed schematic (**c**) of the *demuxalot* algorithm. (1) SNVs are called by aggregating reads from individual transcripts. (2) Base frequencies (**α**) at each SNV location for each droplet barcode are estimated by aggregating called SNVs across transcripts into Bernoulli distributions for each base. (3) Genotype signatures (**β**) at each SNV are initialized from external sources or data as Dirichlet distributions across bases. (4) Genotype assignments (**γ**), represented as categorical distributions for each droplet barcode, are inferred as the likelihood that a given barcode contains a singlet from a given genotype (or doublet from a pair of genotypes). (5) Singlet genotype signatures (**β**) are refined based on assigned genotypes. Steps 4 and 5 define an expectation–maximization (EM) algorithm that can be iterated until genotype signatures and assignments converge. (6) Additional putative SNVs are estimated from the data and added to the base frequencies tensor (columns to the right of the dotted vertical line in **α**). Genotype signatures (**β**) for these new SNVs can also be initialized from data (not shown). Then EM can be rerun to maximally discriminate genotypes.

### Data-efficient SNV calling

Given that individual reads can be noisy due to sequencing, alignment, and PCR duplication errors, SNV calling is inherently error prone. Multiple reads from the same molecule can provide different sources of information about true SNVs at each locus (**Fig. 1c**). Existing methods either incorrectly treat each read as originating from an independent molecule, or ignore additional reads from the same molecule (**Fig. 2a**). In contrast, we maximally exploit the information available from each read by treating SNV calling as a probabilistic estimate of base frequencies at each transcript at each SNV locus. Thus, bases that are highly represented across several reads from the same molecule can be called with high confidence. This method enables calling of up to 3 times more SNV calls per read relative to naive methods (**Fig. 2b**).

**Figure 2.**
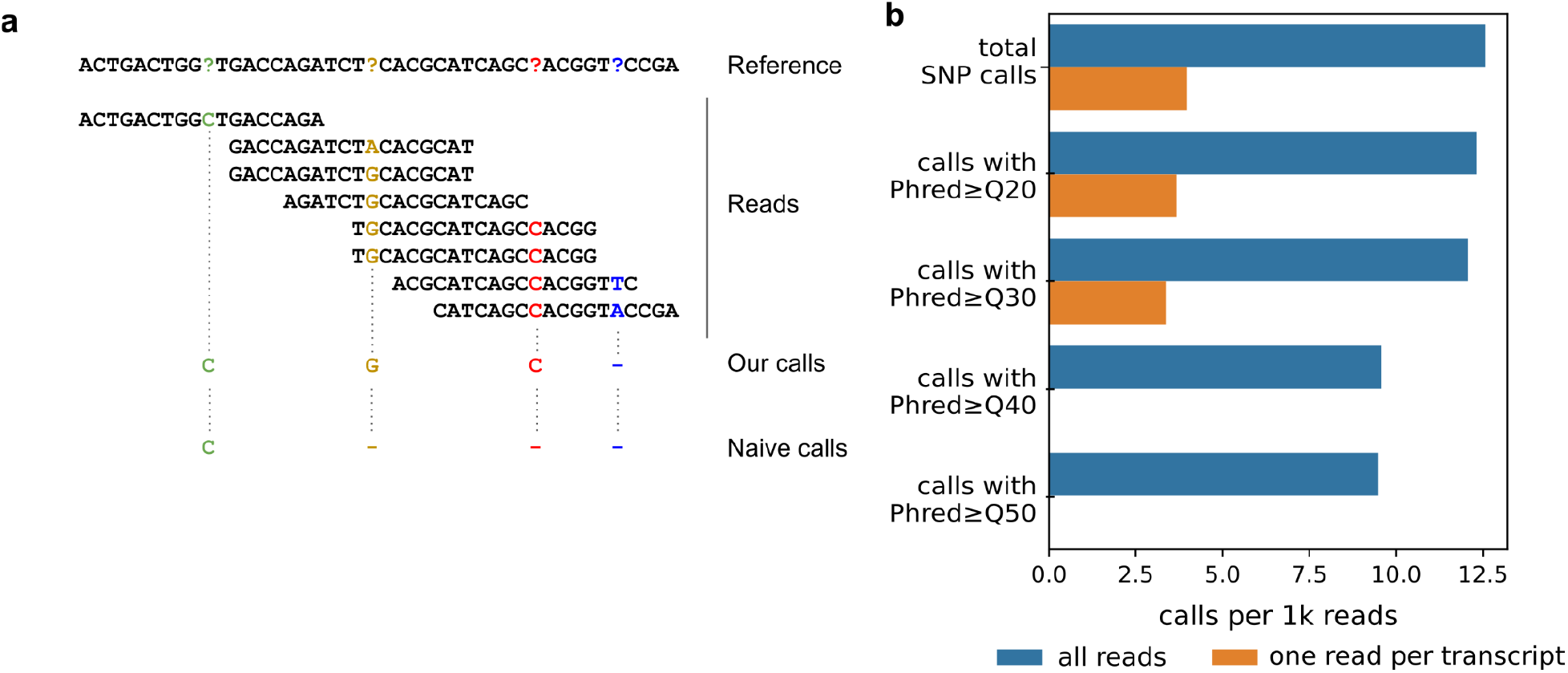
Data-efficient SNV calling. (a) Multiple reads from the same molecule can be used to estimate confidence of each SNV call. SNVs that pass a threshold of confidence are called. (b) Comparison of a naive method that only uses one read per molecule with *demuxalot*’s probabilistic SNV calling that optimally uses information from all reads on data from peripheral blood mononuclear cells (see text). High-confidence calls were supported by multiple reads.

### Flexible and efficient genotype initialization

In multi-donor, multi-batch settings, genotype bead arrays can report different SNVs for different donors depending on choices of technologies or genome references. By maintaining probabilistic estimates of base frequencies for genotype signatures, *demuxalot* enables flexible and data-efficient initialization across heterogeneous data sources. Further, depending on how well donors are represented in common references, certain donors may have very few reference SNVs whereas others may have several. In *demuxalot*, priors for poorly represented donors can be set based on other donors that are well represented.

### Minimal assumptions about allele ratios

In all methods that learn putative SNVs from transcripts, reads are shortlisted based on allele ratios at loci on the whole genome. Most methods in the literature make strongly restrictive assumptions about allele ratios, requiring them to be one of either 0, 0.5, or 1 (Kang et al. 2018; Xu et al. 2019; Heaton et al., 2019; Huang, McCarthy, and Stegle 2019). However there are several scenarios where this assumption fails, including mitochondrial heteroplasmy, trisomy, RNA editing, CNVs (Cooper and Mefford 2011), as well as other scenarios with allele-specific expression (Wang and Valent 2009) such as in X-inactivated cells. *Demuxalot* makes no assumptions about allele ratios, thus generalizing to a wide range of biological scenarios.

*Efficient updates for genotype assignment and refinement*. Many demultiplexing methods, including ours, alternate between genotype assignment and refinement steps (e.g. (Xu et al. 2019; Heaton et al., 2019; Huang, McCarthy, and Stegle 2019). In contrast to other Bayesian methods, *demuxalot* uses tractable models of base frequencies and genotype assignments as conjugate Dirichlet and categorical distributions, which can be updated efficiently in closed form (**see Methods**). In practice, we found that no more than three expectation–maximization (EM) iterations were required for highly accurate genotype assignments for a large number of donors (**Figs. 3, 4**).

**Figure 3.**
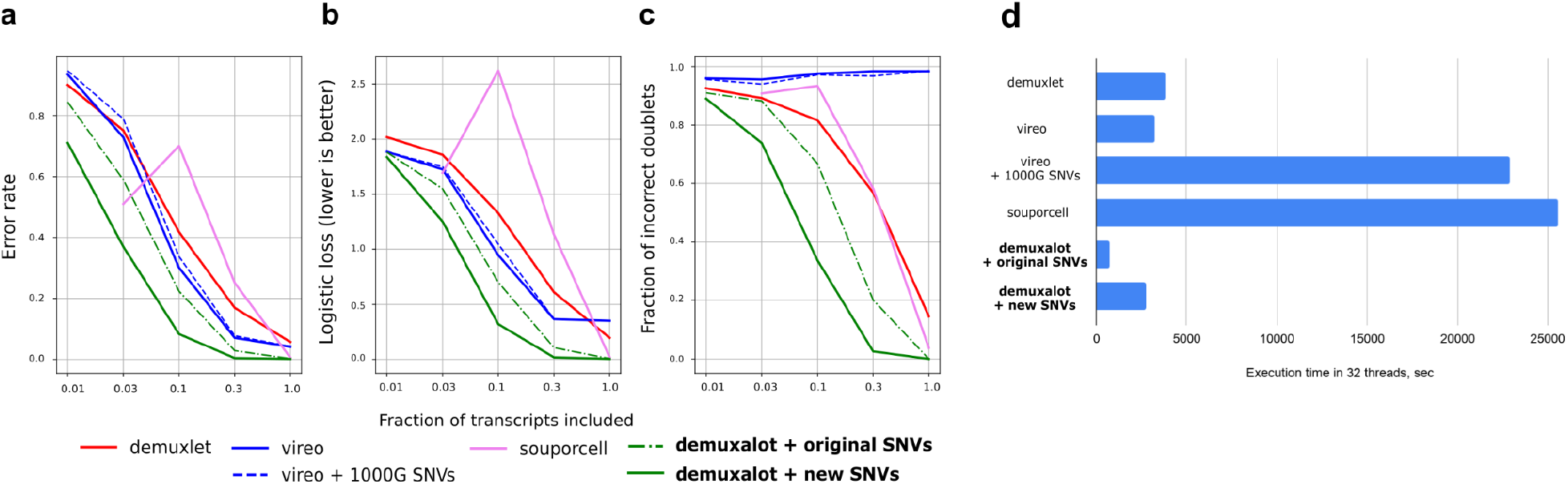
Efficiency of *demuxalot* in the 32-donor setting with a limited amount of transcripts. We use the PBMC dataset from demuxlet paper (see text) and explore the demultiplexing quality of different strategies as a function of the number of available transcripts. Based on knowledge of which genotypes are present in which lanes, we compute (a) fraction of incorrect assignments, (b) logistic loss, and (c) fraction of incorrect doublet assignments among the top 500 doublets predicted by each method. (d) Comparison of runtimes on the full dataset.

**Figure 4.**
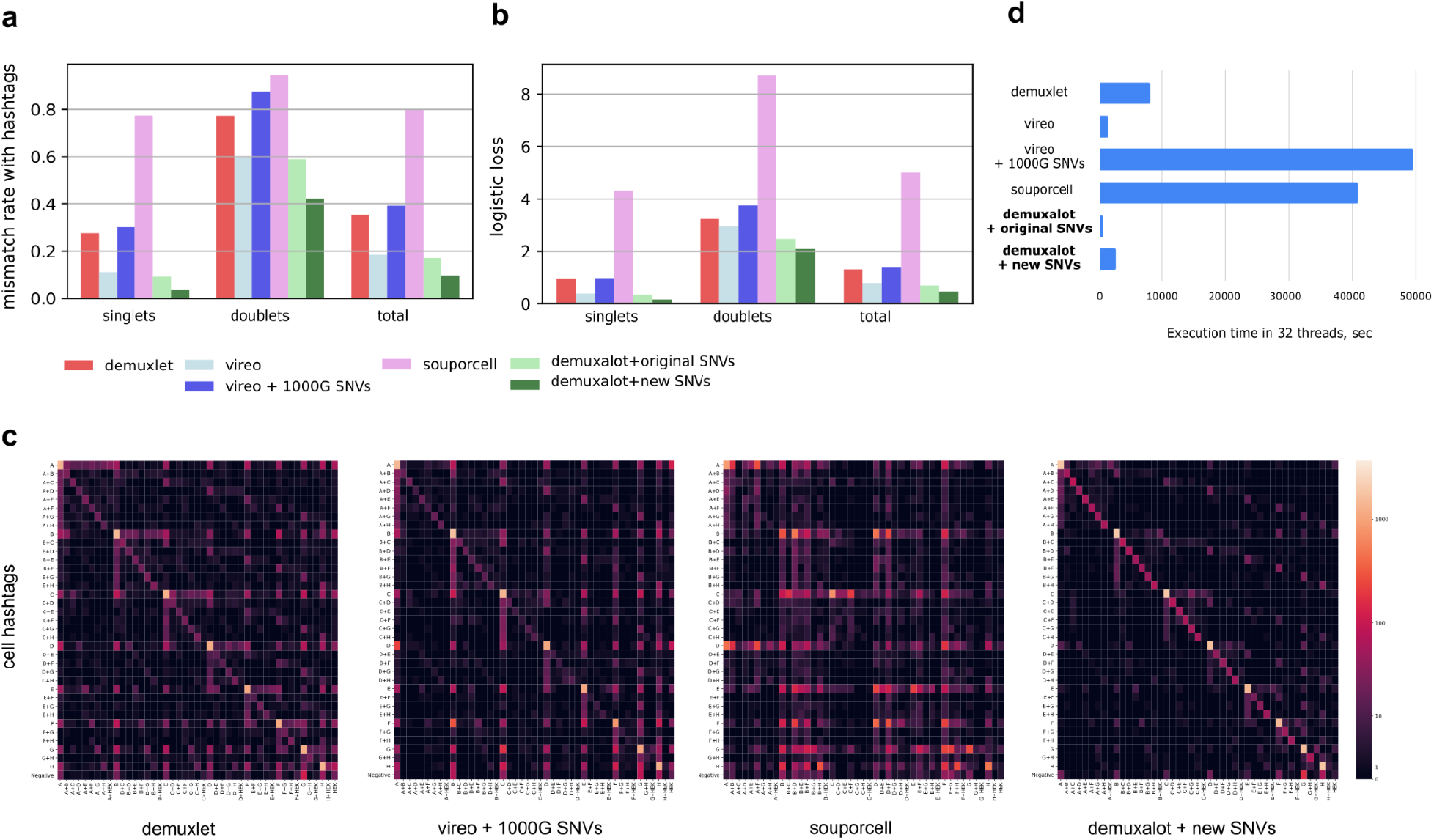
Correspondence between genotype assignments and ground truth sample of origin based on the non-genotype based method of demultiplexing using a cell hashing dataset. For each demultiplexing method, we show the (a) mismatch rate compared with the cell hashing ground truth, (b) logistic loss, (c) confusion matrices (heat maps in log scale), and (d) runtimes.

### Principled framework for SNV enrichment

Several demultiplexing methods enrich the set of SNVs starting from putative SNVs. In contrast to other methods, *demuxalot* only accumulates additional SNVs based on the likelihood that they can distinguish between donors based on the information entropy of base frequency distributions at putative SNV locations (**see Methods**). This strictness makes the method faster and more memory efficient. In the following comparisons we run *demuxalot* with and without SNV enrichment to show contribution of this step.

## More accurate and data-efficient demultiplexing in multi-donor setting

To demonstrate the ability of our approach to deal with a large number of donors, we ran several demultiplexing experiments on the peripheral blood mononuclear cell (PBMC) dataset from the demuxlet paper (Kang et al. 2018). The dataset consists of three lanes, each containing cells from a subset of donors (donors 1-4 in lane A, donors 5-8 in lane B, and donors 1-8 in lane C), which we concatenate while storing information about lane of origin for evaluation. In our experiment, demultiplexers are provided information about 32 genotypes, not just the 8 genotypes in the original study. We evaluated the dataset with the additional genotypes to reflect the use-case of large biobanks when demultiplexing is also used to detect cell line contamination or mishandling during library preparation. Inclusion of additional donors in the set of possible genotype assignments also allows us to measure quality of assignments in the absence of ground truth: for instance, since lane C can contain only cells from donors 1-8, assignment to any other donor is considered incorrect for barcodes from lane C. Similarly doublet predictions are considered incorrect if either of the donors is not from one of the donors in the lane. Since methods can predict drastically different amounts of doublets, doublet detection is evaluated using the 500 most probable barcodes predicted by each method to enable fair comparisons. We found that *demuxalot* results in higher accuracy and lower doublet error rate (**Fig. 3a, 3c**).

To further confirm the stability of this result, we subsampled the library to demonstrate the superior quality of *demuxalot* even when provided with only 30% of transcripts. Due to a more efficient calling procedure, *demuxalot* identifies singlets (**Fig. 3a**) and doublets (**Fig. 3c**) more accurately than other methods for different amounts of available transcripts. Perhaps surprisingly, *demuxalot* shows better performance relative to other methods without SNV enrichment, with enrichment resulting in even further improvement. Complementarily, a lower logistic loss (**Fig. 3b**) confirms well-calibrated assignment probabilities.

We also observed that filtering to common variants (from the 1000 Genomes Project) as suggested by the Vireo authors (Xu et al. 2019; Heaton et al., 2019; Huang, McCarthy, and Stegle 2019) did not improve demultiplexing quality (see Vireo vs Vireo + 1000G in **Figs. 3a-c**). Additionally, we found that souporcell optimization was unreliable, as the quality does not improve with increasing amounts of available transcripts (**Figs. 3a-c**).

In addition to improved accuracy, *demuxalot* achieves faster runtime relative to other methods on the same hardware (**Fig. 3d**). The decrease in runtimes varied from 40% in comparison with demuxlet to 930% in comparison with souporcell, a method that runs expensive realignment of reads as part of the pipeline. A large proportion of this runtime enhancement is attributable to optimized SNV calling procedures (see Methods).

## Better agreement with cell hashing methods

For the PBMC data obtained from the demuxlet paper (Kang et al. 2018), we do not have a ground truth dataset mapping barcodes to samples. Therefore, we used proxy metrics derived from known experimental design parameters to evaluate genotype assignment and doublet detection. For a more rigorous comparison of demultiplexing methods, we sought to further confirm our findings using genotype-independent demultiplexing with cell hashing. In cell hashing, cells are marked with oligo-tagged antibodies prior to pooling. Oligos are sequenced along with RNA and provide ground-truth information about the sample of origin. The cell hashing dataset (Stoeckius et al. 2018) we used contains PBMC samples from 9 donors, 8 of which (A, B, C, D, E, F, G, H) were tagged with antibodies and one (HEK) was left untagged. Thus, a barcode tagged with D can also be a doublet of D+HEK, since HEK cells were not tagged. We address this nuance during accuracy computation.

We computed accuracy (**Fig. 4a**) and logistic loss (**Fig. 4b**) for singlets, doublets, and the full, combined dataset. Compared with existing approaches, *demuxalot* showed superior singlet identification before detecting new SNVs at a fraction of the runtime (**Fig. 4d**). Furthermore, singlet identification significantly improved by including additional putative SNVs detected from the data. Confusion matrices (**Fig. 4c**) further reveal the performance of *demuxalot* relative to other methods. Additionally, we observed that Vireo undercounts doublets by predicting singlets almost always. We also found that souporcell converged to irrelevant but high-confidence assignments, which, in a setting without independent controls, would lead to false confidence in demultiplexing quality. Notably, almost all discordances between our method and cell hashing were doublet vs singlet assignments with one of the singlet genotypes in the doublet (e.g. A+H vs A).

## DISCUSSION

We developed *demuxalot*, a genotype demultiplexing method for single-cell sequencing that is purpose-designed for scaling to large numbers of donors and batches. Using a simple Bayesian model of base frequencies at putative SNVs, we propose a number of improvements to existing methods for SNV calling, genotype initialization and refinement, genotype assignment, and SNV enrichment, which together result in improved computational efficiency and accuracy at scale. Our method’s main contribution to data efficiency comes from the SNV calling step which makes optimal use of multiple reads per transcript. Likewise, its main contribution to computational efficiency comes from dataset-optimized SNV enrichment that aims to maximize genotype information with as few additional putative SNVs as needed. Taken together, these improvements enable scalability to multiple donors and allow for better exploitation of deep sequencing datasets. Finally, the flexible incorporation of genotype priors from multiple sources allows for easy scalability to multiple batches and heterogeneous reference sources.

Beyond algorithmic improvements for data efficiency, accuracy and scale, we also made a number of implementational improvements that lead to higher flexibility and faster runtimes, including the implementation of callbacks to filter out reads, more efficient multithreading over aligned reads, and SNV calling using a single pass through reads. We make the method accessible to users and developers with an open source Python implementation.

With *demuxalot*, we explicitly address shortcomings in published demultiplexing methods: demuxlet, Vireo, scsplit and souporcell (Kang et al. 2018; Xu et al. 2019; Heaton et al., 2019; Huang, McCarthy, and Stegle 2019). Below we highlight areas where *demuxalot* improves upon them. First, unlike all previous methods, *demuxalot* recognizes all reads from the same transcript and optimally estimates the confidence of SNV calls from each such aligned read, resulting in significantly better use of information from deeply sequenced data. Second, demuxlet, the most established demultiplexing method, does not aim to enrich SNVs from the dataset, only relying on genotype priors on reference SNVs obtained from bead arrays (Kang et al. 2018). Like methods that were developed in response to this limitation (Huang, McCarthy, and Stegle 2019; Heaton et al., 2019), *demuxalot* augments information available in genotype priors with putative SNVs from the dataset. Crucially, unlike Vireo or souporcell, our method optimizes this selection of putative SNVs, achieving a superior trade off between computational cost and accuracy than previous methods. Third, Vireo models the exact likelihood of a SNV call using a hierarchical graphical model, and uses approximate variational inference (VI) for parameter estimation. Approximate inference methods like VI that solve a local optimization problem have known weaknesses (Blei, Kucukelbir, and McAuliffe 2017). We found that in practice, Vireo optimization did not always converge to a better solution compared with the initial state. Here, we chose to model the approximate likelihood of SNV calls using a simple conjugate Bayesian model with exact closed-form inference methods for posterior updates, which demonstrates rapid convergence in practice. Fourth, Vireo and demuxlet make limiting assumptions about the prevalence of variant alleles, restricting ploidy to two – i.e., only modeling a reference and an alternate allele. We make no a priori assumptions about allele ratios, implicitly modeling them with Dirichlet distributions over base frequencies at each SNV.

*Demuxalot* achieves very high accuracy in singlet predictions, but doublet identification is still challenging. Although knowing the exact genotype composition of doublets can be valuable to diagnose technical issues in library preparation, it is important to note that for downstream analyses one only needs to discard doublets, not accurately infer their genotypes of origin. In this sense, errors in genotyping doublets are more tolerable than for singlets, but improving doublet accuracy is an important area of future work. In this context, it is worth noting that most demultiplexing papers evaluate doublet detection accuracy using computational simulations, whereas we use data-driven ways to assess quality here. Simulation-based comparisons heavily rely on certain arbitrary assumptions which strongly influence results: e.g., demuxlet quality drops significantly (relative to the original setting) when reads are randomly reassigned to model ambient RNA. Multiple questions arise as to the relevance of these assumptions and if this model truthfully simulates ambient RNA (which one would expect to be a function of cells’ health, which may vary across cultures and depend on cell type or other factors).

A limitation of our method is that it relies on a robust genotype initialization and therefore is not applicable in regimes where no alternate sources of a priori information are available. Although further work could validate our method for de novo demultiplexing contexts, we do not consider de novo demultiplexing to be a viable strategy for large scale applications, where the relative cost of obtaining prior genotype information is vanishingly small.

All pooled single-cell RNA sequencing methods with genetic demultiplexing address the core problems of lane to lane technical variability, high costs per sample, detection and removal of doublets, and the detection and correction of sample mix ups or contaminations. These applications become absolutely essential in multi-donor studies to avoid the introduction of both designed (e.g. lane effects) and accidental (e.g. sample mix ups) confounders. Beyond well-established applications, highly scaled demultiplexing methods like *demuxalot* pave the way for other cutting-edge applications. Modern in vitro systems for disease biology, such as organoids, can be engineered with cells from multiple genetic backgrounds (Jerber et al. 2021) or controlled to express specific alleles such as in isogenic models of X-inactivation (Hinz et al. 2019). For X-inactivation studies, alleles that are active can be distinguished from those that are inactive using the differential information provided by their SNVs. Co-culture systems are becoming increasingly popular in in vitro biology. For applications in neuroimmunology, neuron–astrocyte–microglia tricultures are engineered with primary or iPSC-derived cells from different genetic backgrounds (Guttikonda et al. 2021). Likewise, in vitro patient-derived tumor models often have heterogeneous genetic backgrounds. In each of these cases, single cell sequencing with demultiplexing can computationally isolate cell types and experimental conditions purely via genotype assignments, making it a powerful tool to design and interpret multiplexed experiments at scale. Robust demultiplexing methods that scale to a wide library of genotypes open up possibilities for numerous cutting edge multi-genotype applications in basic science and technology.

## ACKNOWLEDGEMENTS

The authors would like to thank J Sorokin for helpful discussions and comments on the manuscript.

## AUTHOR CONTRIBUTIONS

AR, KS, RB: conceptualization; AR: software development and data analysis; AR and PR: writing—original draft; AR, PR, KS, RB, SK, and GSE: writing—review and editing

## RESOURCES

We have provided an efficient Python implementation of *demuxalot* as an open source library (https://github.com/herophilus/demuxalot).

## METHODS

The basic problem setting is as follows. Each droplet barcode *b* may contain multiple transcripts *t*, tagged with unique molecular identifiers (UMIs). Fragments from each transcript are amplified, fragmented and sequenced as reads *r*. An aligned read may span several SNV positions *s* on the reference genome.

We model the presence or absence of each base (A, T, G, C) at each SNV position *s* on a read *r* spanning that position from transcript *t* and barcode *b* as a categorical distribution Cat(*x*_*s,b,t,r*_).

Calls for transcripts obtained after aggregating over reads are modeled with a categorical distribution Cat(*x*_*s,b,t*_), because transcript can have only one base.

Calls for each barcode *b* are modelled as a set of independent Bernoulli variables, and represent which bases appeared at each SNV position and our confidence in these observations. Specifically, each base call *α*_*s,b,base*_ at SNV position *s* for each barcode *b* is a Bernoulli random variable. There is no restriction on how many of these indicator variables can be turned on at the same time to model scenarios such as trisomy / RNA editing. In this formulation it is of less importance how many transcripts were observed with a particular base called.

We wish to infer a categorical distribution over genotypes for every barcode **γ**_b_, and we model our knowledge about genotype signatures with a Dirichlet distribution *β*_*s,g*_ at every SNV position *s* for every genotype *g*.

We then estimate the probability *p*(**γ**_*b*_ = g) of each barcode *b* being assigned to each genotype *g* in the following sequence of estimation steps.

### Step 1: SNV calling at transcript level

We estimate base frequencies at each SNV position *s* for each transcript *t* for each barcode *b* by aggregating over reads *r*

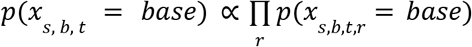

The probability on the right hand side of the equation is computed from the base quality returned by sequencer and the read alignment quality returned by aligner.

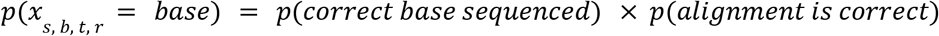

If the probability for the most likely base is 1000 times greater than for any other base (or if only one base is observed in the reads), this base gets assigned and propagated to the following step. Otherwise, no call is made.

### Step 2: SNV calling at barcode level

We estimate base frequencies at each SNV position *s* for each transcript *t* for each barcode *b* by aggregating over transcripts *t*.

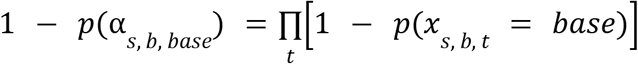

### Step 3: Initialize genotype signature

We set initial genotype signatures (represented as parameters of a Dirichet distribution)

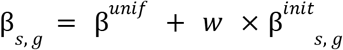

based on GSA array data or other prior information at reference SNV locations *s*. β^*unif*^ guarantees a uniform distribution in the absence of external information (all parameters equal to one).

It is possible to combine several sources of information into the weights. For instance,

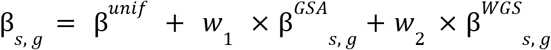

In the absence of information for a given position in reference for genome, we impute β _*s, g*_ using the average over other genotypes. Likewise, when a new SNV is added after the enrichment step (see below), we initialize β with the average over calls for that SNV position *s*._*s, g*_

### Step 4 (E-Step): Genotype assignment

We estimate conditional likelihood of genotype given empirical estimates of barcode base frequencies at SNVs (α_*s, b*_) and current best estimate of genotype signature (β _*s, g*_).

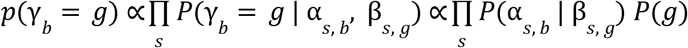

We enrich the set of possible genotypes with “doublet genotypes”, that is genotypes generated by averaging all pairs of two original genotypes. Barcodes attributed to doublet genotypes are subsequently marked as doublets.

Prior probability of genotype *P*(*g*_*singlet*_) = *p* is the same for all genotypes, and *P*(*g*_*doublet*_) = *p* for all doublet genotypes and both are computed from a user-specified expected doublet rate.

### Step 5 (M-Step): Genotype refinement

We marginalize base frequencies over barcodes *b* assigned to genotype *g* for *g* ∈ {1, …, *N*_*g*_} at each SNV position *s*, suppressing the contribution from low-confidence assignments by exponentiating the probabilistic genotype assignment, with exponent q defaulting to 2.

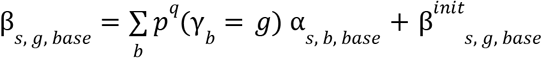

We iterate Steps 4 and 5 until convergence. Empirically, we see convergence in about 3 iterations.

### Step 6: SNV Enrichment

We restrict to barcodes *b* assigned to any one of the genotypes with high confidence *max*_*g*_ *p*(γ_*b*_ = *g*) > *p*_*threshold*_ where default *p*_*threshold*_ is set to 0.7.

For each genotype and position, we collect count statistics of bases *C*_*g, s*_, as follows.

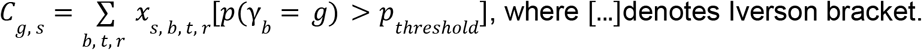

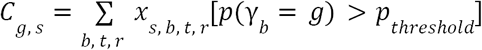, where […]denotes Iverson bracket.

To estimate informativeness *I*of genomic position *s* in identifying genotype *g*, we use

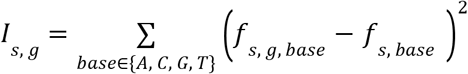

where

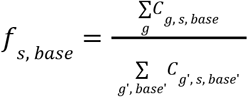

and

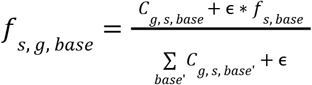

where ϵis a regularizer that anchors the base frequency close to the average across the genotypes if the genotype has a low number of calls for this position.

For every genome, we order variants by information gain *I*_*s, g*_over reference SNVs, select top *k* (default 100) variants.

### Step 7: Iterate steps 4 and 5 until convergence

After one round of SNV enrichment, we reiterate between Steps 4 and 5 (genotype assignment and genotype refinement) until convergence.

### Keeping information about genotypes

We preserve genotypes β_*s, g*_ across runs as opposed to hard assignments (like A/C) or probabilities. This allows storing both observed fraction and confidence individually for each SNV position. Storing the parameters of the Dirichlet distribution affords several advantages.

- Non-informative priors in the absence of data, as well as empirical priors are easily encoded as Dirichlet parameters. In the latter case, these are proportional to the fraction observed in real data without demultiplexing. For example, if A was in 90% of reads at this position, and C in 10%, the empirical prior would be β_*s,g,A*_ = 0. 9, β_*s,g,C*_ = 0. 1
- Fast incorporation of new posterior information from additional scrnaseq runs becomes straightforward.
- Introduction of new genotypes with some missing loci or with some new SNV positions becomes straightforward. This scenario is commonly observed when merging information from different versions of SNV arrays. By imputing low-confidence estimates from other genotypes to missing positions, we can continue to use these positions and still benefit from additional information.
- When the number of genotypes grows, imputation is of high importance, because even if all genotypes are sequenced with the same bead array, there are some positions in each run that the bead array does not “recognize”, each time random. Without this key imputation step, the alternative would result in dropping positions with at least one unknown genotype, which would result in significant loss of information.

### Significant speedup

In our implementation of *demuxalot*, we make a number of algorithmic innovations that contribute to significant speedups. The main contributions to speed are as follows.

- Targeted detection of SNVs reduces the number of positions called. For large biobanks this becomes critical as the number of donor-differentiating positions grows with the number of donors.
- Specifically created calling mechanism with callbacks that makes use of aligner-specific tags to filter alignments. This allows efficient filtering of awkward alignment, similar to a technique previously reported by souporcell ((Xu et al. 2019; Heaton et al., 2019; Huang, McCarthy, and Stegle 2019)) albeit without the considerably expensive realignment step.
- We implement more efficient multithreading by splitting the genome into fragments of variable size to address the common scenario that some fragments have much deeper coverage and thus should take more time to process.
- Calling is made with a single pass though the alignments, not by observing SNV-after-SNV as in other tools. This is slower for small lists of SNVs, however it becomes drastically more efficient as the number of SNVs grows and a single read overlaps more than one SNV.

